# AuroraB-kinase methylation by SETD6 regulates cytokinesis and protects cells from chromosomal instability

**DOI:** 10.1101/2024.12.22.629973

**Authors:** Michal Feldman, Anand Chopra, Dikla Nachmias, Kyle K. Biggar, Daniel Sevilla, Natalie Elia, Dan Levy

## Abstract

SETD6 is a non-histone lysine methylatransferase, previously shown to participate in several housekeeping signaling pathways such as the NFkB pathway, Wnt signaling pathway, mitosis and more. In the current study we show evidence that SETD6 methylation is involved in the regulation of cytokinesis - the final process that divides cell contents into two daughter cells. SETD6 depleted HeLa cells presented high levels of chromatin bridges and actin patches, which are commonly observed following chromosomal segregation errors. In a proteomic screen we identified Aurora-B as a novel SETD6 substrate. Aurora-B kinase is an essential regulator of cytokinesis, known to actively delay cytokinesis as a response to the presence of chromatin in the midzone. We found that SETD6 binds and methylates Aurora-B on two adjacent lysine residues. Upon replication stress, Aurora-B methylation by SETD6 increases but is abolished when the two lysine methylation targets are substituted. In addition, replication stress led to a high tendency of SETD6 depleted cells to multinucleate, a major chromosomal-instability (CIN) phenotype. We detected a significant reduction in the Aurora-B kinase activity during cytokinesis in SETD6 knockout cells upon replication stress, which could be the mechanism underlying the accumulation of CIN phenotypes in these cells. CIN is a hallmark of cancer and is associated with tumor cell malignancy. Our findings suggest that Aurora-B methylation by SETD6 carries meaningful implications on tumorigenic cellular pathways.

## INTRODUCTION

Non-histone protein methylation has been shown to take a notable part in the post translation modifications network, with many implications in the regulation of diverse cellular pathways, playing a role both in physiological and disease related processes^1^. One of the most basic cellular pathways, regulated by non-histone lysine methylation, is the cell-cycle which ends with the division of the cell into two daughter cells^2–5^. An accurate completion of the cell division process is of crucial importance for the cellular ability to function and preserve its genetic load properly and therefore this process represents a hallmark in cancer development^6^. The cell cycle includes several checkpoints, which are surveillance mechanisms that monitor the order, integrity, and fidelity of the major events of this process^7^. Proteins that regulate these checkpoints are thus attractive biomarkers as well as therapeutic targets for inhibition of tumor initiation and progression. Among these checkpoints, the one determining the accurate timing of daughter cells separation is the NoCut checkpoint, which occurs at the final step of mitosis - cytokinesis^8^. The NoCut checkpoint delays abscission in response to chromosome segregation defects and is dependent on the protein Aurora-B, located at the spindle midzone between the two daughter cells. Aurora-B belongs to a family of serine/threonine kinases, all considered essential kinases for cell division, each participating in a different mitotic step and localized at its unique cellular position ^9^. Aurora-B starts at early G2 phase and localizes to the chromosomes in prophase, the centromere in prometaphase and metaphase, the central spindle in anaphase and the midbody in cytokinesis^10^. During cytokinesis Aurora-B senses the presence of chromatin within the bridge, indicating a problem in DNA segregation. Such chromatin aberrant structures are often the consequence of replication stress events that occurred during S (DNA synthesis) phase, and their presence might lead to delayed abscission time, regulated by Aurora-B^11^. In the current study we observed that cells lacking the methyltransferase SETD6, demonstrated aberrant chromatin segregation within the intercellular bridge. This observation was consistent with the detection of specific methylation of Aurora-B by SETD6 on two adjacent lysines. Moreover, in the absence of SETD6, cells presented a high tendency to multinucleate when replication stress was induced. We discovered that Aurora-B methylation maintains its kinase activity, which is essential for its role as a chromatin segregation sensor. These results imply that Aurora-B methylation by SETD6 protects cells from gaining different segregation phenotypes, shortly described as Chromosomal Instability (CIN). Chromosomal instability (CIN) is a characteristic of many human cancers, and it is associated with poor prognosis, metastasis, and therapeutic resistance. CIN results from errors in chromosome segregation during mitosis leading to structural and numerical chromosomal abnormalities^12^. Investigating the mechanism through which SETD6-mediated methylation of Aurora-B regulates CIN accumulation, will uncover a novel path in neoplastic transformation and label this specific methylation event as an suitable target for cancer therapy.

## RESULTS

### Depletion of SETD6 leads to the presence of segregation aberrations during cytokinesis

We were interested to find out whether SETD6 is involved in more advanced steps of mitosis, such as anaphase and cytokinesis, critical processes which determine chromosomal segregation. To test the effect of SETD6 on chromosomal integrity during late mitotic stages, we stained control and SETD6 KO cells with DAPI (DNA staining) and LaminA/C, a nuclear lamina component which has a major role in chromatin binding and organization^13^ (Fig. 1A). Surprisingly, we discovered that a significant portion of all captured cytokinetic cells presented chromatin bridges (reflected by LaminA/C staining) upon SETD6 depletion (Fig. 1A). To establish this finding, we captured actin-filaments patterns in these cells using phalloidin staining. In cytokinesis with chromatin bridges, cells delay abscission and retain actin patches at the intercellular canal to prevent chromosome breakage ^14^. Actin patches were indeed observed at a higher rate in the SETD6 KO HeLa cells, compared to control cells (Fig. 1B). Chromatin bridges and actin patches are known to reflect segregation defects. SETD6 is known to localize mainly to the cell nuclei, however this information was obtained from interphase cells ^15^. The possible involvement of SETD6 in cytokinesis led to the hypothesis that SETD6 might localize adjacent to the intercellular bridge during cytokinesis. Using ectopic expression of GFP-SETD6, we detected SETD6 in the middle of the tubulin bridge (Fig. 1C), supporting its relevance to the cytokinesis and abscission processes.

**Figure 1.**
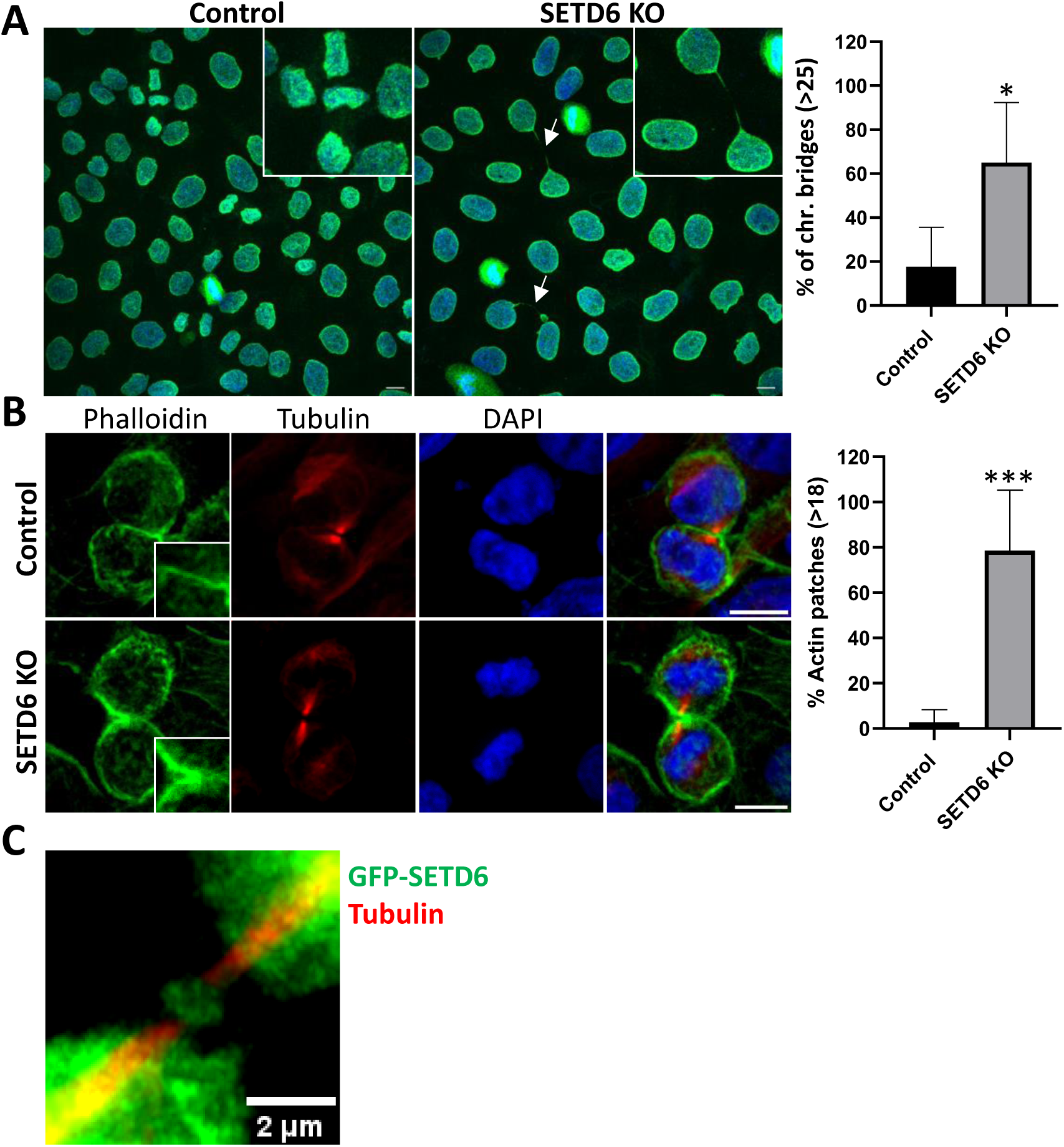
Depletion of SETD6 leads to the presence of segregation aberrations during cytokinesis. Control and SETD6 KO HeLa cells were fixed and stained with anti-LaminA/C Ab and DAPI to detect chromatin bridges (A) or with anti-Tubulin Ab, Phalloidin and DAPI to visualize actin patches (B) The percentage of cells presenting chromatin bridges or actin patches was calculated as the fraction of these cells out of all cytokinetic cells observed in the field of view. Statistical analysis was performed using the unpaired, two-tailed *t* test. *: *P* < 0.01, ***: *P* < 0.0001. Scale, 10μM. (C) Control HeLa cells were transfected with GFP-SETD6 for 48 H, then fixed and stained with anti-Tubulin Ab to visualize SETD6 within the cytokinetic bridge. Scale, 2μM.

### SETD6 binds Aurora-B and methylates it

The cellular difficulty to complete proper cytokinesis, as observed in Figure 1, implied for the existence of a novel SETD6 substrate, such that is involved in regulating cytokinesis and chromosomal segregation. In a proteomic screen, in which more than 9500 proteins were tested, we identified 118 SETD6 substrates, out of which more than 25% are associated with oncogenic signaling pathways ^16,17^. One of the identified substrates was the Polo-like kinase 1 (PLK1) protein, which was later validated as a SETD6 substrate ^5^. Methylation of PLK1 by SETD6 was shown to maintain proper mitotic pace, significantly during the early mitotic events: prometaphase and metaphase ^5^. One of the substrates that were revealed in the screen mentioned above, was the Serine/Threonine kinase Aurora-B (Fig. 2A). Aurora-B kinase (from now on referred to as: AurB) is a part of the Aurora kinases family. It commences to express at early G2 and localizes to the chromosomes in prophase, the centromere in prometaphase and metaphase, the central spindle in anaphase and the midbody in cytokinesis ^10^. During cytokinesis AurB is located at the spindle midzone and functions as the NoCut checkpoint regulator – it senses the presence of chromatin within the bridge, indicating a problem in DNA segregation to the two daughter cells ^8^. Such chromatin bridges are often the consequence of replication stress events that occurred during the S phase, and their presence might lead to delayed abscission time, regulated by AurB ^11^. To validate the methylation of AurB by SETD6, we performed an *in-vitro* methylation assay in the presence of purified recombinant GST-SETD6 and His-AurB, using ^3^H-labeled S-adenosyl methionine as the methyl donor. As shown in Fig. 2B, we found that His-AurB is directly methylated *in-vitro* by GST-SETD6. AurB-SETD6 interaction in cells was examined using immunoprecipitation of HA-SETD6 and Flag-AurB over-expressed in HeLa cells and western blot analysis. HA-SETD6 was found to precipitate together with Flag-AurB, implying that these two proteins associate in HeLa cells (Fig. 2C). Since AurB is mostly functional during mitosis, its SETD6-mediated methylation was tested in cells during mitosis. Control and SETD6 knockout HeLa cells ^5^ were synchronized to G1/S phase using double-thymidine block. Cells were lysed 8 hours after thymidine release (which coincides with mitosis), and immunoprecipitation was performed using a pan-methyl antibody. Western blot for AurB revealed a significant reduction in methylation signal in SETD6 depleted cells (Fig. 2D). Overall, these results demonstrate and confirm that AurB is a methylation substrate of SETD6 during mitosis.

**Figure 2.**
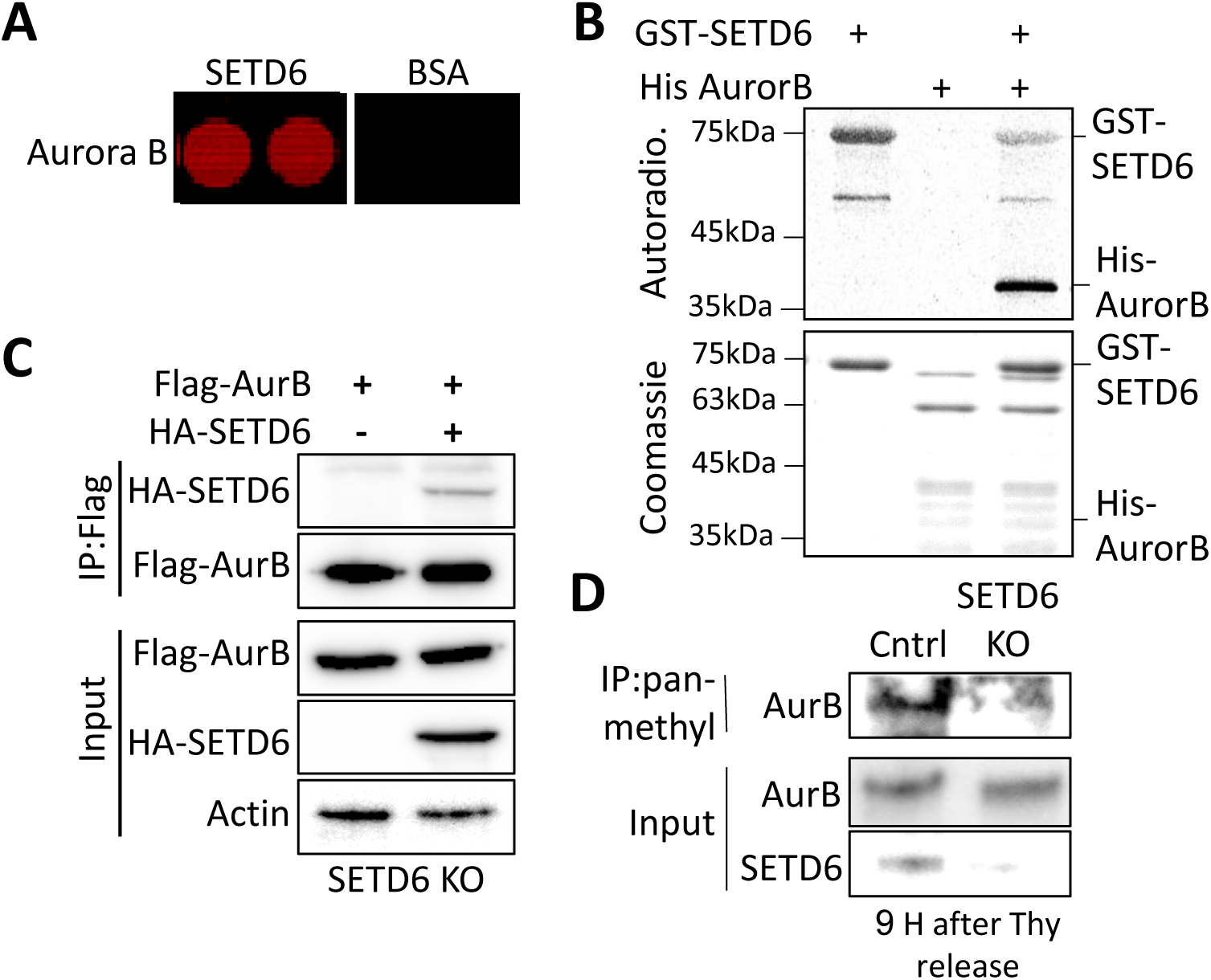
SETD6 methylates AurB and interacts with it. (A) Positive methylation signal of recombinant AurB by GST-SETD6 shown in a Protoarray experiment. Protoarrays were incubated over-night under PKMT reaction conditions with GST-SETD6 and GST as a negative control. Arrays were then probed with a pan-methyl antibody followed by a fluorophore-conjugated secondary antibody. (B) *In vitro* methylation assay in the presence of ^3^H-labeled S-adenosyl methionine with recombinant His-AurB and GST-SETD6, incubated together overnight. Coomassie stain of the recombinant proteins used in the reactions is shown on the bottom. (C) SETD6 KO cells over-expressing Flag-AurB with or without HA-SETD6 were subjected to Immunoprecipitation with Flag magnetic beads and blotted against AurB and SETD6 to test their interaction. (D) Control and SETD6 KO HeLa cells were synchronized using double thymidine block and harvested 9 H after block release. Methylated proteins were immunoprecipitated from lysates and blotted against AurB.

### In the absence of SETD6, cells tend to multinucleate upon replication stress

As the master regulator of the NoCUT checkpoint, AurB senses the pathological presence of mis-segregated DNA in the cytokinetic bridge and while delaying the recruitment of the abscission complex, it stabilizes a canal between the two daughter cells to prevent chromosome breakage^18^. To study the role of AurB methylation by SETD6 during cytokinesis, we induced a mild replication stress in control and SETD6 depleted HeLa cells using minor amounts of HydroxyUrea or Aphidicolin as previously indicated ^19^. Such replication stress would not arrest the cells through the NoCUT checkpoint but would force the cells to delay the abscission to avoid chromosomal damage. Since delaying abscission is under the responsibility of AurB, we stained these cells with tubulin to image the intercellular bridges and DAPI. Images captured and quantified showed a significant increase in the percentage of the cytokinetic cells in the control cells upon replication stress, indicating a delayed abscission (Fig. S1). Such an increase was not detected in the SETD6 KO cells, pointing out a possible deficiency in the cytokinesis checkpoint. While observing these cells, we noticed an elevation in the percentage of multinuclear cells in the SETD6 KO cells upon replication stress induction, in comparison to the control cells (Fig. 3A and 3B). Using live imaging, we detected cells demonstrating a regression of the cleavage furrow during cytokinesis, leading to cytokinetic failure and to tetraploidization (Supporting video 1). We next aimed to link the severe phenotype detected in SETD6 depleted cells under replication stress (Fig. 3A) to the physiological SETD6-mediated methylation level of AurB. When cells were immunoprecipitated using a pan-methyl antibody, an increased methylation signal was observed in control cells following Aphidicolin treatment, a signal which almost completely disappeared in the absence of SETD6 (Fig. 3C). These findings imply that SETD6-mediated methylation of AurB is involved in the abscission regulation. This involvement appears to be more emphasized during replication stress, when the necessity for a fully active NoCUT checkpoint machinery rises.

**Figure 3.**
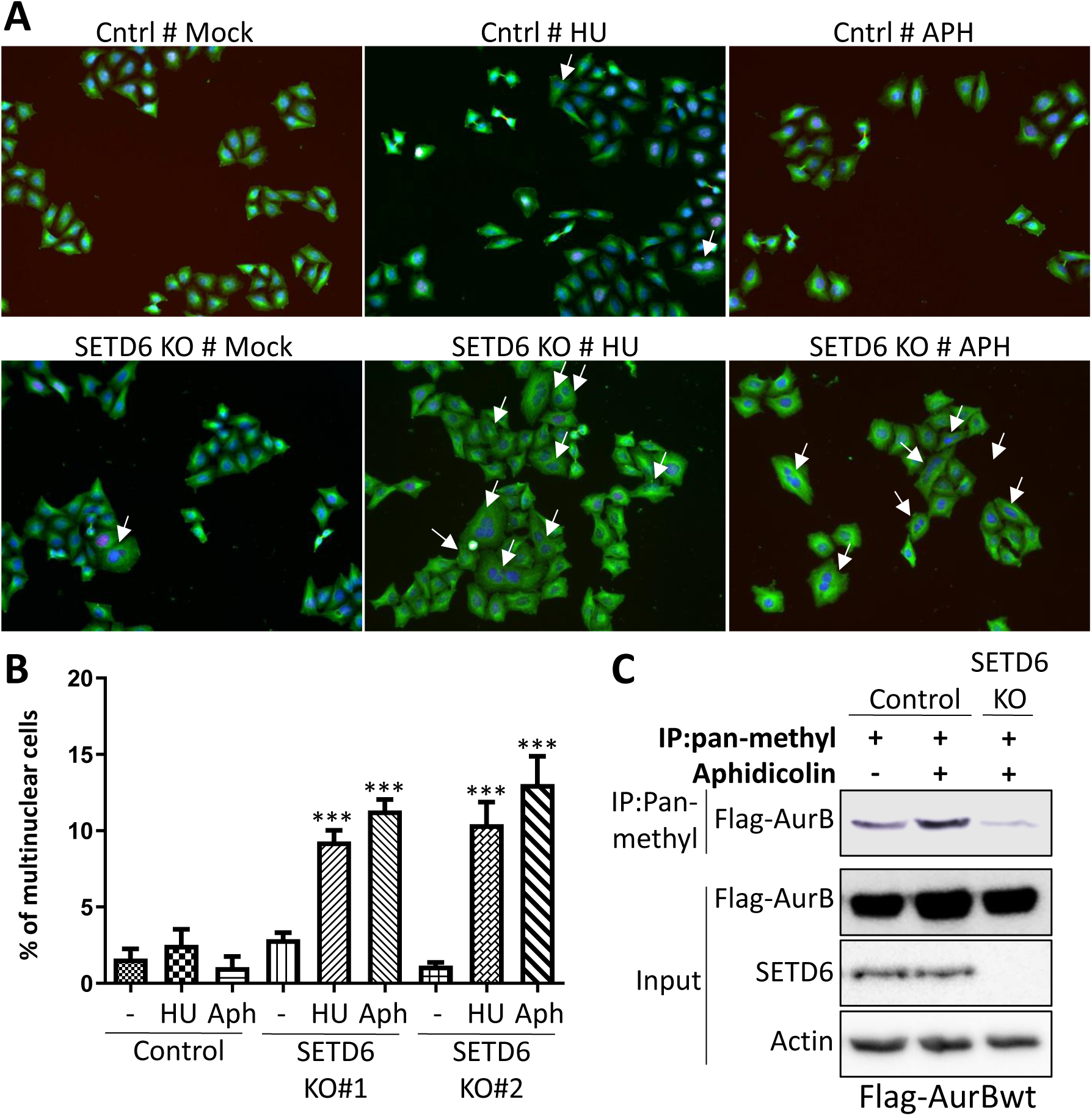
AurB methylation by SETD6 is required during replication stress. (A) Control and SETD6 KO HeLa were treated or not with 50 μM HydroxyUrea (HU) or 0.4 μM Aphidicolin (APH) for 24 H, after which cells were fixed and stained with Anti-Tubulin Ab and DAPI. Representative images are shown of 20 images captured for each condition. Multinuclear cells are indicated in white arrows. (B) Multinuclear cells were manually counted in control and two different SETD6 KO clones. Statistical analysis was performed using the unpaired, two-tailed *t* test. ns, not significant. ***: *P* < 0.0001. (C) Control and SETD6 KO HeLa cells, were transfected with Flag-AurB for 48 H and treated or not with 0.4 μM Aphidicolin for 24 H. lysates from these cells were subjected to immunoprecipitation using pan-methyl Ab and blotted against Flag-AurB.

### SETD6 methylates AurB on lysines 194&195

To specifically link AurB methylation by SETD6 to AurB cellular activity, we aimed to map the methylation site. To find AurB methylation site, we performed a radioactive peptide array in the presence of SETD6 and all AurB lysine residues containing peptides as wildtype and with K-R substitution (Fig. 4A). These experiments revealed that SETD6 methylates AurB at two adjacent lysine residues (K194 and K195). The methylation of these residues was validated in cells by immunoprecipitating methylated proteins overexpressing wild-type FLAG-AurB, single and double mutants harboring mutations in K194 and K195. As shown in Figure 3B, we observed a slight decrease with each of the single mutants K194R and K195 (see band quantification under the gel) compared to wild-type AurB. Interestingly, an 80% decrease in signal intensity was observed in the double mutant, suggesting that K194 and K195 are the primary sites in AurB targeted for methylation by SETD6. Consistent with these results, we confirmed by mass spectrometry experiments that AurB methylation on both K194 and K195 is SETD6-dependent, as depletion of SETD6 significantly reduced the methylation signal whereas rescue with HA-SETD6 rescued the signal back (Fig. 4C). To examine the physiological relevance of this methylation event to cytokinesis regulation, we ectopically expressed Flag-AurB wt and K194&195R double mutant in HeLa cells and tested cytokinetic aberrations upon replication stress. A significant increase in chromatin bridges, detected with Lamin A/C staining (Fig. 4D) and actin patches, detected with phalloidin staining (Fig. 4E) were observed upon over-expression of AurBK194&195R mutant. These findings confirmed that the specific methylation of AurB by SETD6 protects cells from accumulating segregations aberrations upon replication stress.

**Figure 4.**
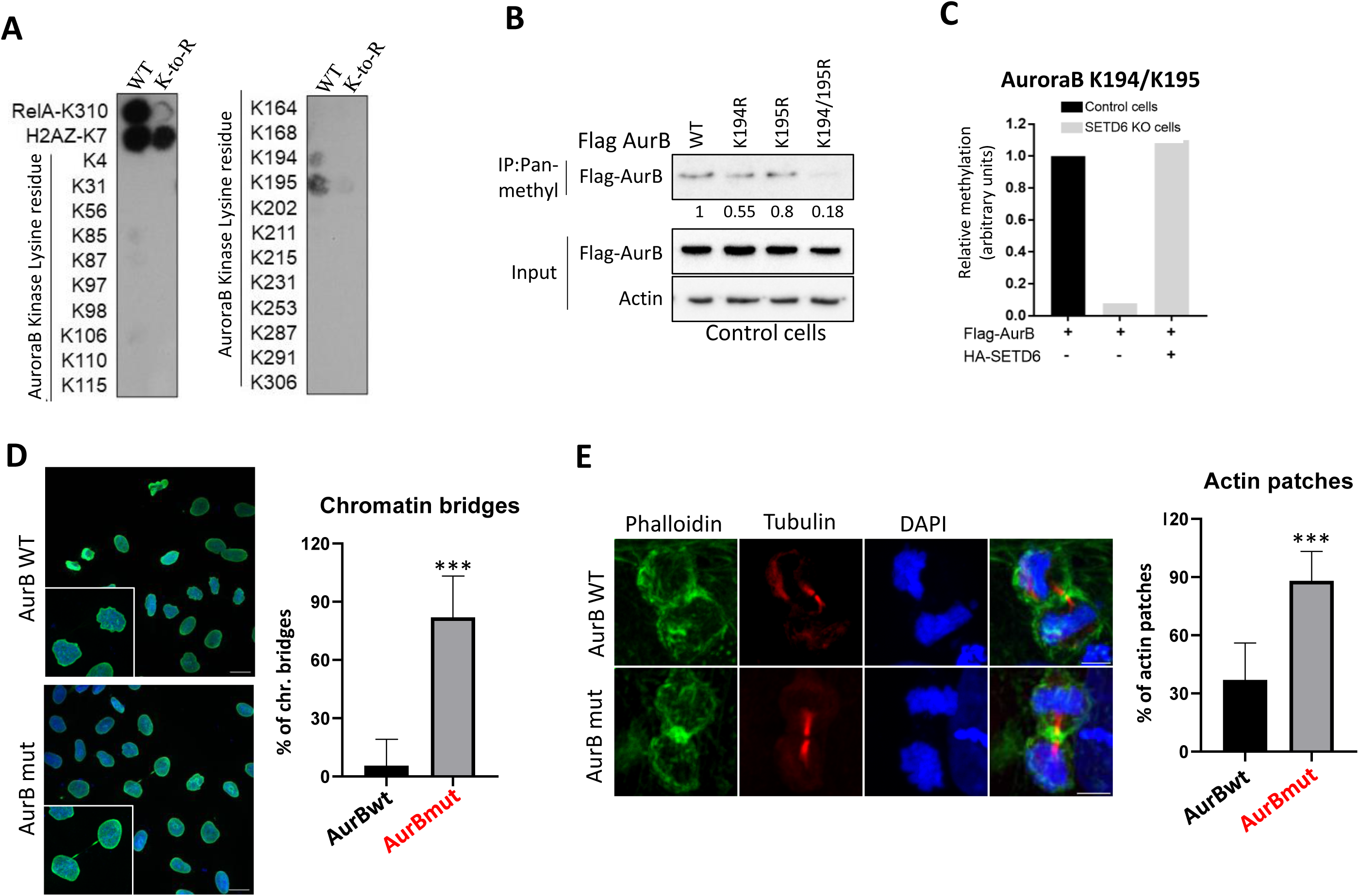
SETD6 methylates AurB on lysines 194&195. (A) Radioactive peptide array of all AurB lysine residues containing peptides as WT and with K-R substitution. Peptides containing RelA-K310 and H2AZ-K7 were used as positive controls. B. Hela control cells were transfected with either Flag-AurB_WT_, Flag-AurB_K194R_, Flag-AurB_K195R_ (single mutants) or Flag-AurB_K194_195R_ (double mutant) expressing plasmids. (B) Methylated proteins were immunoprecipitated from these cellular extracts with a pan-methyl antibody and were then blotted against Flag-AurB. (C) Quantification of methylated AurB_K194_195R_ by mass spectrometry in control (black bar), SETD6 KO cells transfected with empty or HA-SETD6 (grey bars). Methylation of AurB_K194_195R_ mutant decreases in comparison to AurB_WT_. (D+E) Control and HeLa cells were transfected with Flag-AurB_WT_ or Flag-AurB_K194_195R_ for 48 H, then fixed and stained with anti-LaminA/C Ab and DAPI to detect chromatin bridges (D) or with anti-Tubulin Ab, Phalloidin and DAPI to visualize actin patches (E) The percentage of cells presenting chromatin bridges or actin patches was calculated as the fraction of these cells out of all cytokinetic cells observed in the field of view. Statistical analysis was performed using the unpaired, two-tailed *t* test. \*\*\**P* < 0.0001. Scale, 10μM.

### AurB activity is dependent on its methylation following replication stress

AurB is known for its role as the NoCUT checkpoint master regulator during cytokinesis. Cytokinesis in animal cells depends on the completion of chromosome segregation, which is necessary to prevent CIN features which could lead to severe defects such as multinucleation by furrow regression. Most cells with chromosome bridges suppress furrow regression and continue to proliferate normally ^11^. Unsegregated chromatin in the cleavage plane is usually sensed by the NoCut checkpoint regulators to control abscission timing and to protect mis-segregating cells against tetraploidization by furrow regression ^11^. In order to sense the presence of chromatin in the intercellular bridge and further act to delay abscission during cytokinesis, AurB needs to be accurately localized at the spindle midzone, where it responds to DNA segregation errors^8^. We therefore hypothesized that SETD6-mediated methylation of AurB might have an effect on AurB position along the intercellular bridge. Using structured-illumination-microscopy (SIM), we observed no significant methylation-dependent changes in the midbody position of AurB, either when looking at control and SETD6 KO cells, under replication stress or not (Fig. S2A) or when over-expressing AurBwt versus AurBK194_195R mutant (Fig. S2B). These observations led to the conclusion that lack of methylation by SETD6 does not lead to the accumulation of CIN phenotypes by modulating AurB position. Another significant requirement for AurB activity within the midzone is its phosphorylation state: Auto-phosphorylation on threonine 232 located at the kinase domain is essential for the integrity of its kinase activity ^20^. Under normal conditions, levels of phosphorylated T232-AurB in the midbody of cytokinetic cells are reduced to allow abscission. However, AurB maintains high phospho-activity at the intercellular bridge when mis-segregated chromosomes are sensed within the bridge ^19,21^. Interestingly, lysines 194&195 are located within the kinase domain of AurB, required for the phosphorylation of its substrates (Fig. S3). To examine whether methylation of AurB has an impact on its phospho-activity under replication stress, we used control and SETD6 KO cells in which replication stress was induced or not using Aphidicolin. These cells were stained with DAPI (nuclear staining), anti-tubulin antibody and an antibody detecting the T232 phospho-AurB (Fig. 5A). Fluorescence intensity was quantified and a reduction in the T232 phospho-AurB signal was recorded upon SETD6 depletion with or without aphidicolin (Fig. 5B). Staining the cells with the replication stress marker 53BP1^19^, showed a similar increase in its signal between the control and SETD6 KO cells upon Aphidicolin treatment (Fig. S4A and S4B), indicating that replication stress was induced in a SETD6 methylation-independent manner. 53BP1 is a DNA damage response protein which protects vulnerable lesions following replication stress until repair can occur ^22^. The observation that 53BP1 is induced in SETD6 KO cells upon replication stress suggests that CIN phenotypes are not a result of defects in the stress sensing cellular machinery during G1 and S phases. Instead, it supports our notion that the NoCUT checkpoint is malfunctioning in the absence of SETD6 methylation of AurB. This observation was confirmed by detecting a decrease in phospho-AurB fluorescence in cells stably expressing Flag-AurBK194_195R mutant in comparison to the wild-type protein (Fig. 5C-5E). These findings indicate that methylation of AurB by SETD6 regulates AurB kinase activity by affecting its phosphorylation state. These findings could explain why depletion of SETD6 leads to CIN characteristics.

**Figure 5.**
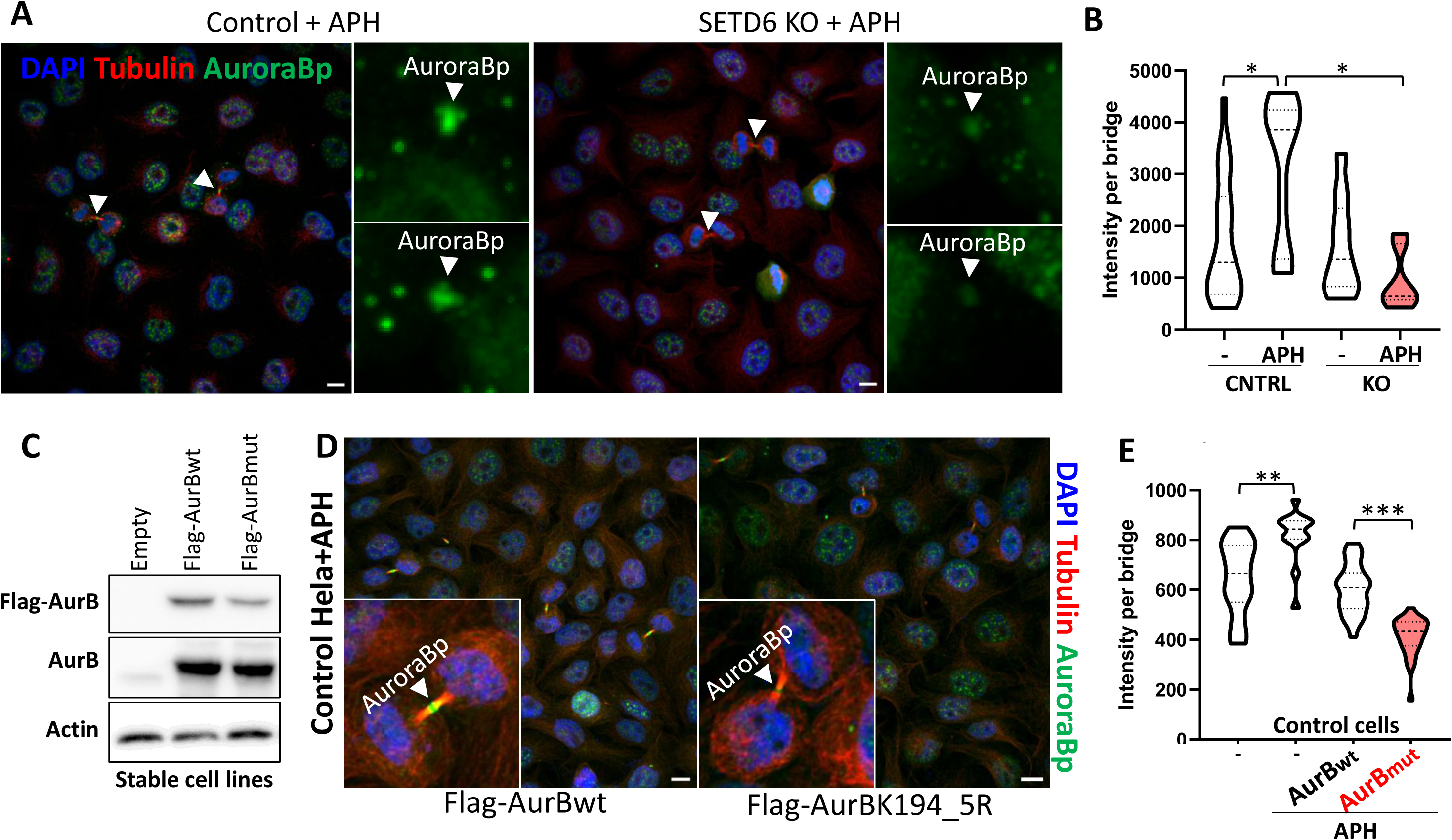
Lack of methylation leads to reduction in AuroraB phospho-activation. (A) Control and SETD6 KO HeLa cells treated or not with 0.4 µM Aphidicolin, were fixed and stained with anti-Tubulin antibody (Red), and anti-AurB-pT232 (green). (B) Quantification of fluorescence intensity of midbody oriented AurB-pT232 staining (*n* ≥ 7 cells per condition). (C) Western blot analysis of Flag-AurB_WT_ and Flag-AurB_K194_195R_ stable expression in HeLa control cells. (D) Similar staining procedure was performed in HeLa control cells stably expressing Flag-AurB_WT_ or Flag-AurB_K194_195R_ treated or not with 0.4 µM Aphidicolin. (E) Quantification of fluorescence intensity of midbody oriented AurB-pT232 staining (*n* ≥ 14 cells per condition). Cells were visualized by a confocal microscope (63×). Magnification of the midbody area is shown for each image. Statistical analysis was performed using the unpaired, two-tailed *t* test. \**P* < 0.01, **: *P* < 0.01, ***: *P* < 0.0001. Scale, 10μM.

### Depletion of AurB methylation affects its ability to recruit downstream partners required for abscission

The integrity of AurB kinase performance is essential for recruiting its substrates to the midbody, a necessary event for activating the NoCUT downstream signaling pathway, resulting in abscission^18^. AurB-dependent phosphorylation of the midbody ring component MKLP1 and the ESCRT-III component CHMP4C is considered vital for abscission delay^23^. Since the decrease in phosphor-active state of AurB is known to reflect its ability to bind and phosphorylate substrates, we assumed that methylation deprivation might sequentially lead to the loss of this ability. Using co-immunoprecipitation, we found that AurBwt bound MKLP1 and CHMP4C to a higher extent when replication stress was induced (Fig. 6A, 2nd and 3rd lanes). This binding was abolished when AurB could not be methylated (when the AurBK194_195R mutant was used for IP in control cells or when the AurBwt was used for IP in the SETD6 KO cells, Fig. 6A, lanes 4 and 5, respectively). A similar reduction in binding to unmethylated AurB was also shown for the ATPase VPS4, which is the downstream substrate of the ESCRT machinery, also required for timing the abscission event^23^.

**Figure 6.**
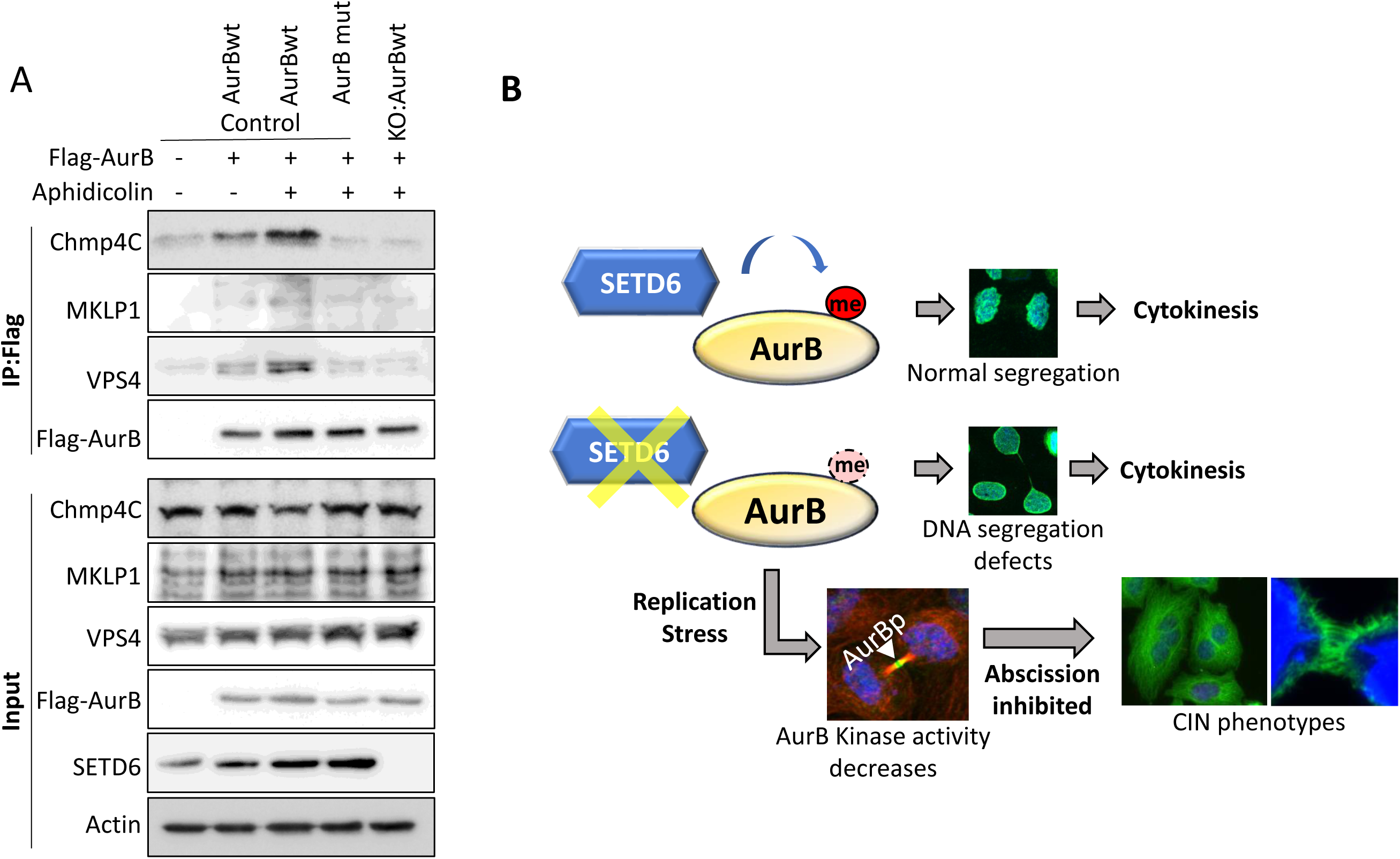
AurB interaction with its downstream partners decreases when AurB is not methylated. (A) Control and SETD6 KO HeLa cells over-expressing or not Flag-AurB_WT_ or Flag-AurB_K194_195R_ mutant were treated or not with o.4 uM Aphidicolin for 24 H. Lysates were subjected to immunoprecipitation with Flag magnetic beads and blotted against the indicated antibodies. (B) Schematic model describing the effect of SETD6 methylation on AurB auto-activity during replication stress. When methylation is abolished, AurB kinase activity decreases, leading to the accumulation of CIN phenotypes within the cells.

Our results strongly suggest that AurB methylation by SETD6 protects cells from gaining CIN features by maintaining AurB performance during cytokinesis following replication stress.

## DISCUSSION

Numerous studies in the past few years have shown evidence establishing the significant role of non-histone lysine methyltransferases in the regulation of different steps of the mitotic process, among them: SETD7/9^2^, G9A^4^ and SETD6^5^. These reports highlight the trending understanding that lysine methylation is an integral part of the machinery required for a well-orchestrated cell division process. Moreover, it was recently reported that the methyltransferase SMYD2 regulates cytokinesis through lysine methylation of the ESCRIII component CHMP2B^3^. In the absence of this methylation, abscission was shown to be delayed due to modulation of the ESCRTIII complex dynamics ^3^. In the current study we revealed that the master NoCUT checkpoint regulator AurB is directly and specifically methylated by the methyltransferase SETD6. AurB was shown to be hyper-methylated by SETD6 upon replication stress induction. Following the dysregulation of AurB lysine methylation, either through loss or aberrant gain, AurB auto-phosphorylation was compromised at the midbody zone. This directly affected its kinase activity within the ICB and modulated its binding affinity for various substrates, eventually leading to the accumulation of severe CIN characteristics (Fig. 6B). Chromosomal instability (CIN) is a hallmark of many cancer cells, contributing to tumor progression and therapeutic resistance. We demonstrate that the methylation of AurB by SETD6 plays a significant role in preventing CIN phenotypes, highlighting a novel layer of regulation in mitotic control that is crucial for cell division integrity. Our findings underscore the significance of SETD6-mediated AurB lysine methylation during cytokinesis, specifically how this modification regulates its kinase activity and interaction with substrates crucial for cytokinetic completion. Altered methylation of AurB during this phase can result in defects in cleavage furrow formation, modulated abscission, and overall failure of cytokinesis, leading to multinucleation and aneuploidy, hallmark features of CIN^24^. This study demonstrates that AurB methylation is essential for the proper resolution of cell division, ensuring that chromosomal segregation is completed without errors. These findings not only enhance our understanding of the molecular mechanisms underlying CIN, but also highlight potential therapeutic strategies for targeting AurB methylation in cancer.

## MATERIALS AND METHODS

### Cell lines, transient and stable transfection, replication stress and Thymidine block

HeLa cells and Human embryonic kidney cells (HEK293T) were maintained in Dulbecco’s modified Eagle’s medium (Sigma) with 10% fetal bovine serum (Gibco), 2mg/ml L-glutamine (Sigma), penicillin-streptomycin (Sigma), and non-essential amino acids (Sigma). Cells were cultured at 37°C in humidified incubator with 5% CO_2_. Transfection was carried out using *Trans*IT^®^-LT1 (Mirus) or Lipofectamine 2000 (Life Technologies) reagents, according to manufacturer’s instructions. Transfection was performed for 24-48 h before harvesting or fixating the cells. For stable transfection, HEK293T were transfected with pLENTI-empty or pLENTI-FLAG-AurB_WT_ or pLENTI-FLAG-AurB_K194_195R_ with the plasmids encoding the VSV and gag-pol. Control or SETD6 KO HeLa cells were infected with the produced viruses and selected on 500ug/ml Hygromycin. For replication stress induction, 0.4 μM Aphidicolin or 50 μM HydroxyUrea were added to the cells for 24 H, after which cells were harvested for western blotting analysis or fixed for Immunofluorescence analysis. For double-Thymidine block, HeLa cells were plated at 30% confluence, then supplemented with 2mM Thymidine (Sigma, T1895-1G) for 17 H. After which, cells were washed once with PBS, and fresh media was added to the cells for 9H. Cells were then supplemented for additional 15 H with 2mM Thymidine, after which cells were washed once with PBS, and fresh media was added. Following Thymidine release, cells were harvested at indicated times and analyzed by Western blot.

### Antibodies, Immunoprecipitation and Western blot analysis

Antibodies used: anti-AurB (Abcam, ab2254), anti-AurBpT232 (Rockland, 600-401-677S), anti-pan-methyl (Abcam, ab23366), anti-SETD6 (Genetex, GTX629891), anti-Flag M2 (Sigma, F1804), anti-HA (Millipore, 05-904), anti-Actin (Abcam, ab3280), anti-LaminA/C (Invitrogen, MA5-35284), anti-tubulin (Abcam, ab6046), MKLP1 (Santa-Cruz, sc-867), VPS4 (Sigma, SAB4200025), Chmp4C (biorbyt, orb326595) and 53BP1 (Abcam, 175933). HRP-conjugated secondary antibodies (goat anti-rabbit and goat anti-mouse) were purchased from Jackson ImmunoResearch (111-035-144 and 115-035-062, respectively). For western blot analysis, cells were homogenized and lysed in RIPA buffer (50mM Tris-HCl pH 8, 150mM NaCl, 1% NP-40, 0.5% sodium deoxycholate, 0.1% SDS, 1mM DTT and 1:100 protease inhibitor cocktail (Sigma). Samples were heated at 95°C for 5min in Laemmli sample buffer and resolved by SDS-PAGE, followed by western blot analysis. Immunoprecipitation was performed using anti-FLAG M2 magnetic beads (Sigma, M8823) or A/G agarose beads (Santa Cruz, SC-2003) conjugated to anti-pan-methyl. Briefly, ∼200µg of proteins extracted from cells using RIPA buffer were incubated over-night at 4⁰C with 15µl pre-washed FLAG M2 magnetic beads or pan-methyl conjugated A/G beads. The beads were then washed 3 times with RIPA buffer, heated for 5min in Laemmli sample buffer at 95⁰C and resolved on 7%-10% SDS-PAGE gel followed by western blot analysis.

### Generation of AurB lysine substitutions

Site directed mutagenesis for the generation of AurB lysine mutants were amplified using the following primers, followed by DNA sequencing for confirmation, and cloned into pCDNA3.1-Flag (for cell ectopic expression) and pLENTI-Flag (for lentiviral stable transfection of cells). For the AurB K194R substitution the following primers were used: forward 5’-TAATGTACTGCCATGGGcgcAAGGTGATTCACAGAGACATAAAGC-3, reverse 5’-GTCTCTGTGAATCACCTTgcgCCCATGGCAGTACATTAGAGC-3’.

For the AurB K195R substitution the following primers were used: forward 5’-ATGTACTGCCATGGGAAGcgcGTGATTCACAGAGACATAAAGCCAG-3’, reverse 5’-TATGTCTCTGTGAATCACgcgCTTCCCATGGCAGTACATTAGAGC-3’.

For the double AurB K194_195R substitution the following primers were used: forward 5’-CTAATGTACTGCCATGGGcgccgcGTGATTCACAGAGACATAAAGCCAG-3’, reverse 5’-TATGTCTCTGTGAATCACgcgcgcCCCATGGCAGTACATTAGAGC-3’.

### Immunofluorescence

For Immunofluorescence (IF) fluorescently labeled secondary antibodies were used: Alexa488 anti-mouse (Invitrogen, R37120), Alexa488 anti-rabbit (Invitrogen, A21441) Alexa647 anti-mouse (Invitrogen, A21463) and Alexa594 anti-rabbit (Abcam, ab150080) and Phalloidin staining (Invitrogen, O7466). For immunofluorescence (IF), cells were fixed with 4% PFA, permeabilized with 0.5% Triton X-100 and blocked with 10% fetal bovine serum for 30 min, after which cells were stained for 3 h with the relevant primary antibody (described above) followed by 30 min incubation with fluorescent secondary antibody and DAPI. For the quantifications of multinuclear cells and cytokinetic cells, slides were analyzed using the EVOS FL cell imaging station (ThermoFisher). For fluorescence quantification, images were acquired with 3i Marianas (Denver, CO) spinning disk confocal microscope using a 100x or 63x Zeiss Plan-Apochromat oil, 1.4 NA objectives. Each frame represents maximum intensity projection for Z-stacks (0.27μm) captured. Analysis of images was performed using Fiji as described below.

### Image analysis

For the quantification of mean fluorescence intensity of phospho-AurB, the borders of tubulin bridges (using tubulin staining) were manually segmented in all stacks of each field using the wand tracing tool in the fiji software. Integral fluorescence density for corresponding channel was calculated within the defined bridge area and normalized to the bridge volume. Similarly, 53BP1 fluorescence intensity was measured within the nuclei, segmented according to DAPI segmentation.

### Recombinant protein expression and purification

*Escherichia coli* BL21 transformed with a plasmid expressing His-AurB were grown in LB media. Bacteria were induced by IPTG induction and homogenized with ice-cold lysis buffer containing phosphate-buffered saline (PBS), 10mM imidazole, 0.1% Triton X-100, 1mM PMSF and 0.25mg/ml lysozyme for 30 min. The lysates were then subjected to sonication on ice (18% amplitude, 1 min total, 10 sec ON/OFF) and purified on a His-Trap column using AKTA Pure protein purification system (GE) by elution with 0.5M imidazole in PBS buffer, followed by overnight dialysis (PBS, 10% glycerol). Recombinant SETD6 was over-expressed and purified from insect cells as previously described ^25^.

### *In vitro* methylation assay

Methylation assays were performed with recombinant proteins. Methylation reaction (total volume of 25µl) contained 4µg His-AurB and/or 4µg GST-SETD6, 2mCi ^3^H-labeled S-adenosyl-methionine (Perkin-Elmer) and PKMT buffer (20mM Tris-HCl pH 8, 10% Glycerol, 20mM KCl, 5mM MgCl2). The reaction tube was incubated over-night at 30°C. Methylation reactions were resolved by SDS-PAGE for Coomassie staining (Expedeon, Instant*Blue*) and exposed to autoradiogram.

### Peptide array

Peptide spot methylation assay was performed as previously described^26^. 0.77 uM His-SETD6 was incubated with the array overnight at room temperature in 50 mM Tris pH 8, 20 mM KCl, 5 mM MgCl2, and 10% glycerol. The reaction was performed in 3 mL containing 8 uL ^3^H-labeled S-adenosyl-methionine (Perkin-Elmer) and the array contained 48 peptides spots at 2 nmol each.

### Mass spectrometry analysis

Control and SETD6 KO HeLa cells were transfected with Flag-AurB_WT_, with or without HA-SETD6 for 48 H. Cells were then harvested and lysates were subjected to overnight immunoprecipitation using Flag magnetic beads. Immunoprecipitated proteins were alkylated and then digested on-beads with AspNI. Resulting peptides were then analysed by targeted mass-spectrometry.

### Live cell imaging

Cells were plated at low density in four-well chamber slides (Ibidi), transfected with GFP-tubulin and mCherry-H2B and imaged 24 H later. Z stacks of cells were collected for 12 hours at 10 minutes intervals using a fully incubated confocal spinning-disk microscope (Marianas; Intelligent Imaging, Denver, CO) with 40×oil objective (numerical aperture, 1.3) and were video recorded on an EMCCD camera (Evolve; Photometrics, Tucson, AZ). Image processing and analysis were done using SlideBook version 6 (3I Inc.).

### Structured illumination microscopy (SIM)

Image acquisition and processing was performed as previously described^27^. In short, cells were plated at low density on high-resolution #1.5 coverslips (Marienfeld, Lauda-Konigshfen, Germany) and fixed using 4% PFA. Cells were further subjected to immunostaining as described above. 3D SIM imaging of low expressing cells was performed using the ELYRA PS.1 microscope (Carl Zeiss MicroImaging). Thin z-sections (0.11 to 0.15 μm) of high-resolution images were collected in 3 rotations and 5 phases for each channel. Image reconstruction and processing were performed in ZEN (Carl Zeiss MicroImaging).

### Statistical analysis

Statistical analyses for all assays were performed with GraphPad Prism software, using one-way, two-way analysis of variance (ANOVA) or student’s t-test.

## Acknowledgments

We thank the Levy lab for technical assistance and helpful discussions. This work was supported by grants to DL from The Israel Science Foundation (262/18 and 496/23), The Israeli Cancer Research Foundation Israel (ICRF), and from the Israel Cancer Association.

## Author Contribution

MF, and DL conceived and designed the experiments. MF performed the majority of the experiments. AC and KKB performed the mass-spectrometry and peptide arrays experiments. DN, DS and NE assisted in the live imaging microscopy experiments and data analysis. MF and DL wrote the paper. All authors read and approved the final manuscript.

---

**Figure S1.**
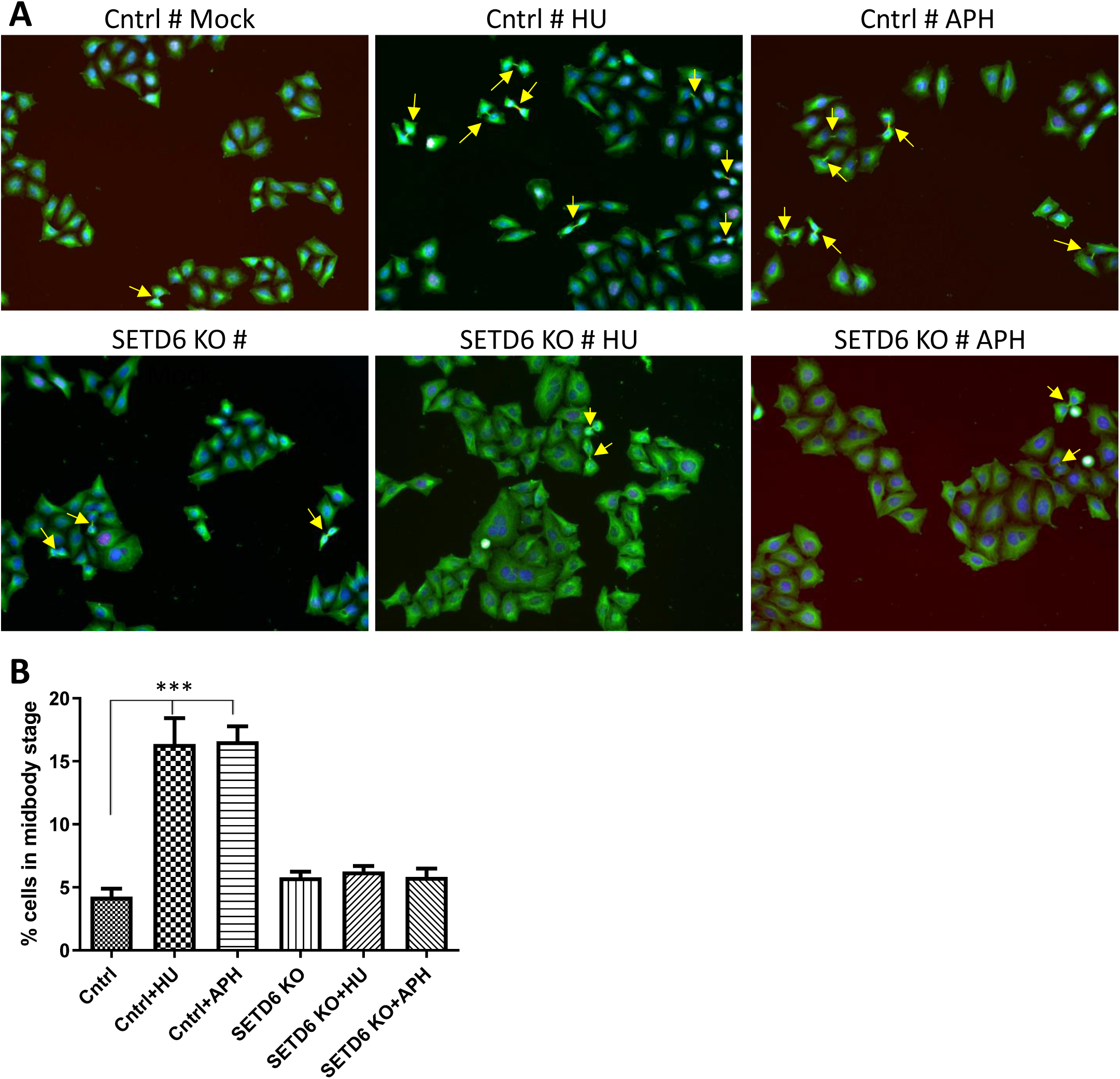
Cytokinesis is delayed in control cells upon replication stress, but not in SETD6 KO cells. (A) Control and SETD6 KO HeLa cells were treated or not with 50 μM HydroxyUrea (HU) or 0.4 μM Aphidicolin (APH) for 24 H, after which cells were fixed and stained with Anti-Tubulin Ab and DAPI. Representative images are shown of 20 images captured from each condition. (B) Cytokinetic cells were manually counted in control and two different SETD6 KO clones. Statistical analysis was performed using the unpaired, two-tailed *t* test. \*\*\**P* < 0.0001.

**Figure S2.**
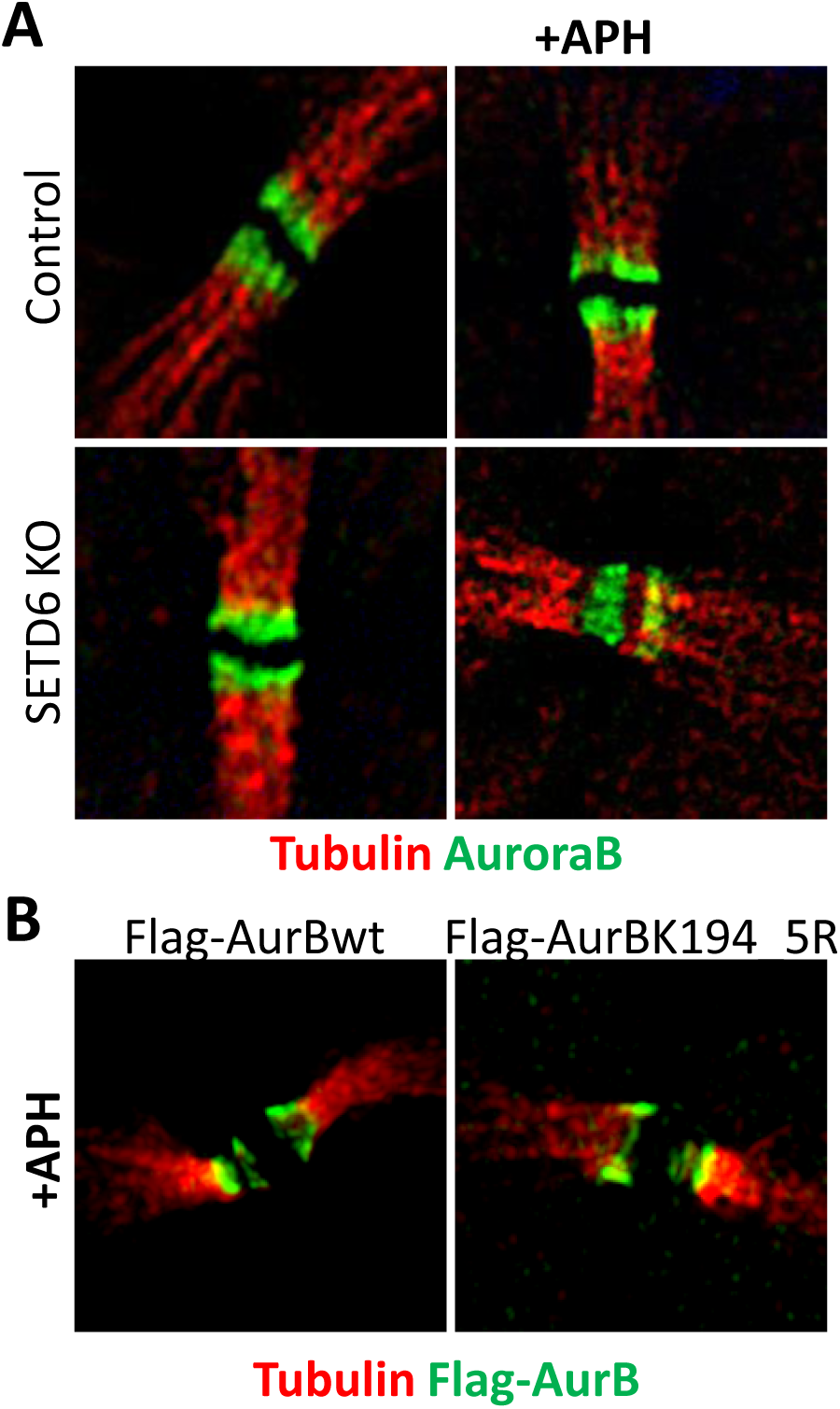
AurB localizes to the intercellular bridge when methylated or not. SIM imaging shows normal AurB localization in control & SETD6 KO HeLa cells (A) and in HeLa control cells over-expressing Flag-AurB_WT_ or Flag-AurB_K194_ mutant (B). Cells were treated or not with 0.4 μM Aphidicolin for 24 H, fixed and stained with the indicated Antibodies.

**Figure S3.**
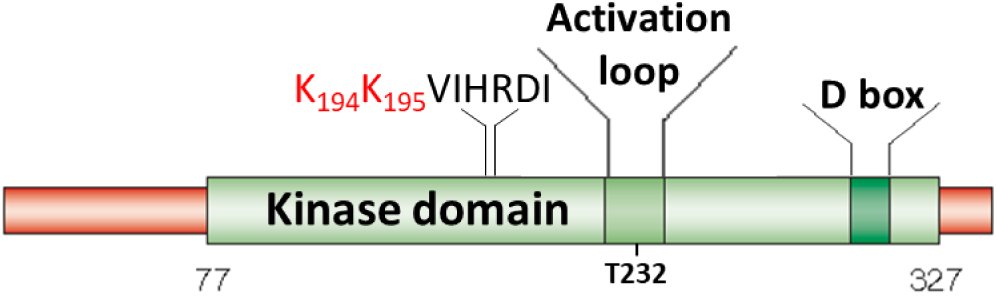
Schematic illustration of AurB domain structures. AurB protein contains the kinase domain, destruction box (D box) domain and activation loop. The localization of the SETD6-methylated Lys-194&195 residues is indicated, as well as the Thr-232 residue, which is located within the activation loop and is auto-phophorylation is essential for AurB kinase activity.

**Figure S4.**
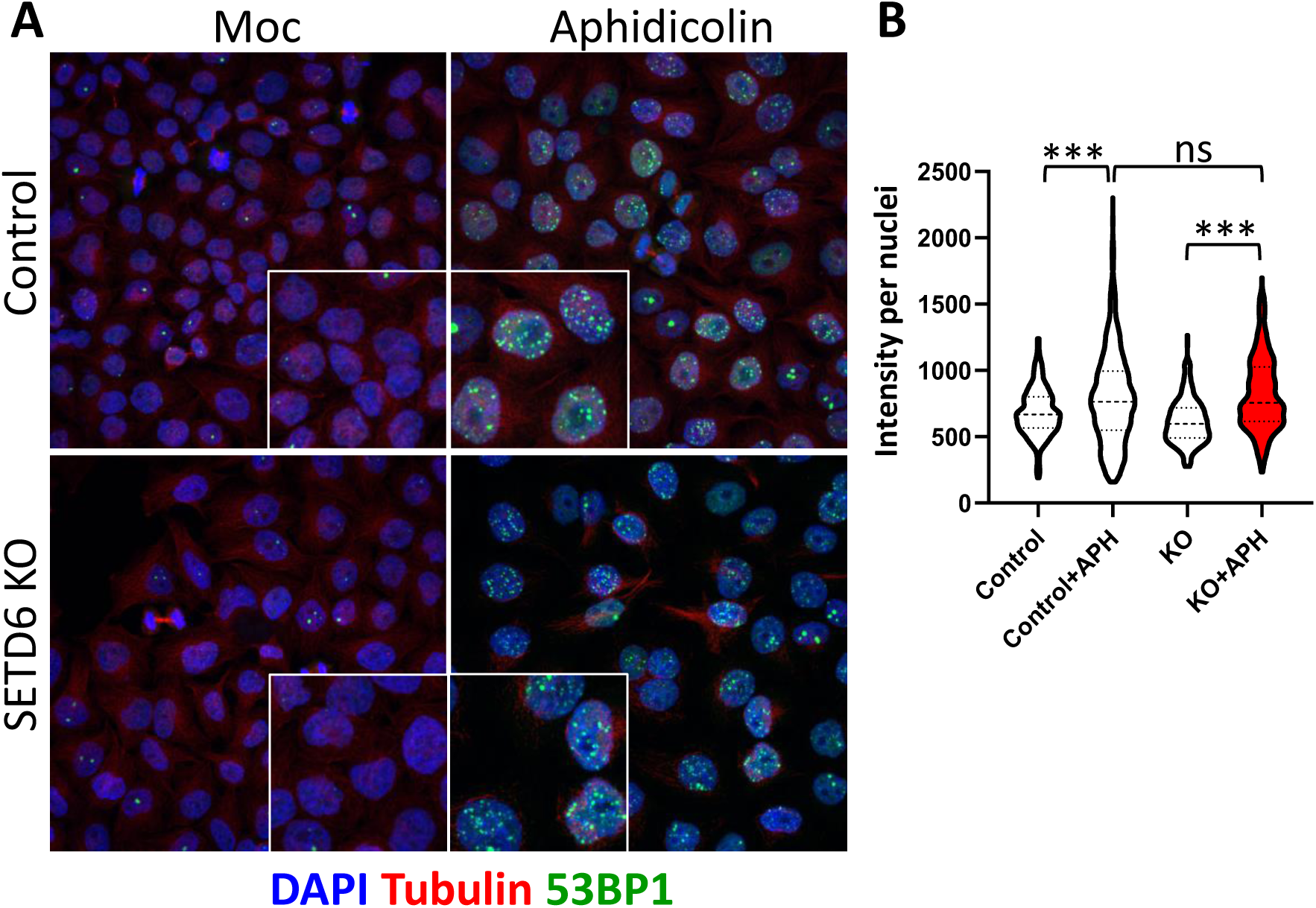
Lack of methylation does not affect cellular replication stress response. (A) Control and SETD6 KO HeLa cells treated or not with 0.4 µM Aphidicolin, were fixed and stained with anti-Tubulin antibody (Red) and anti-53BP1 antibody (green). (B) Quantification of fluorescence intensity of 53BP1 staining within the nuclei. Cells were visualized by a confocal microscope (63×). Magnification of the nuclei area is shown for each image. Statistical analysis was performed using the unpaired, two-tailed *t* test. ns=not significant, ***: *P* < 0.0001.

**Supplementary video 1.**
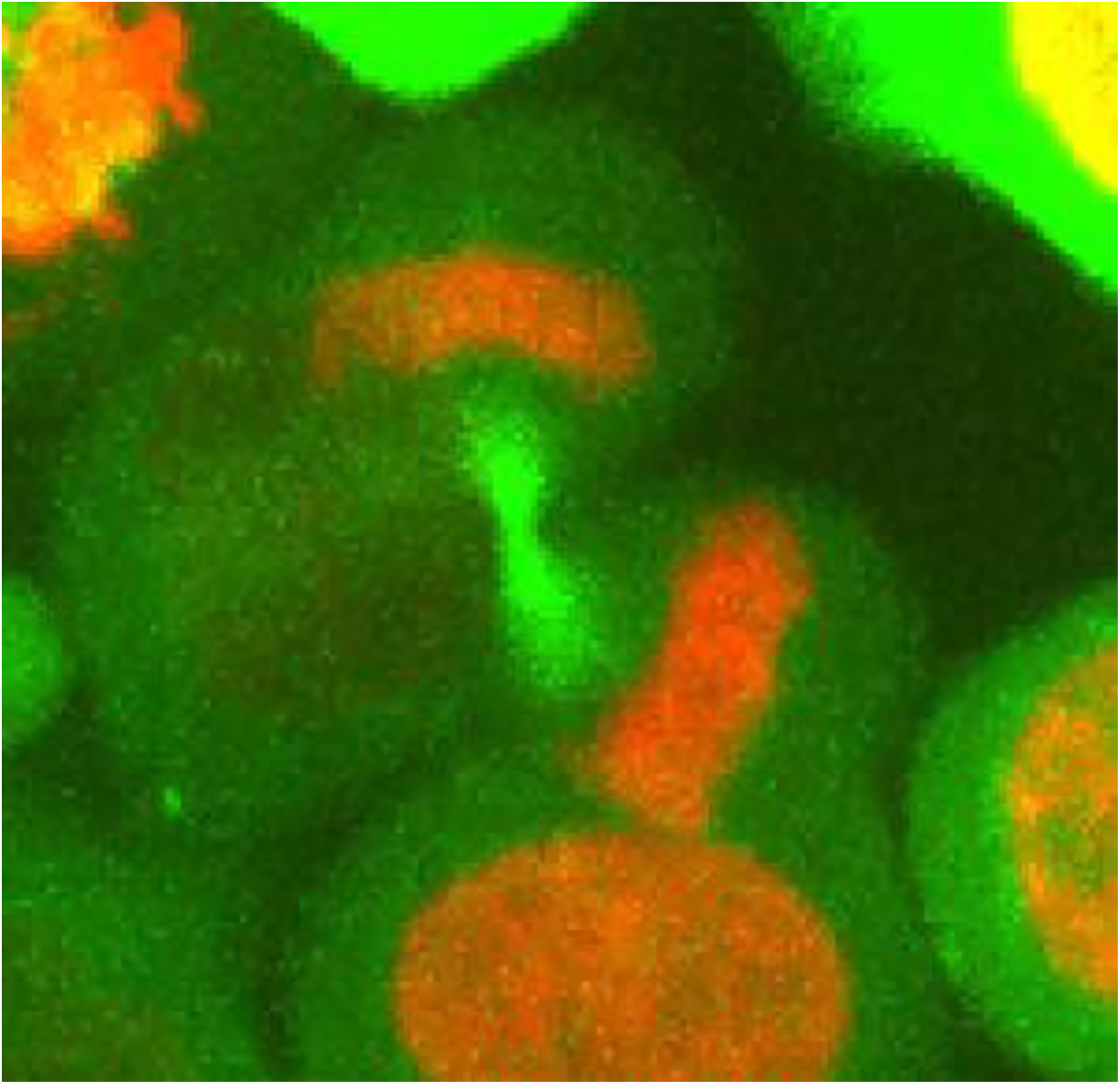
Depletion of SETD6 leads to multinucleation upon replication stress induction. SETD6 KO HeLa cells expressing Tubulin-GFP and mCherry-H2B were submitted to live imaging using a confocal spinning-disk microscope.

